# Identification of a suitable substrate for the growth and development of *Hibiscus sabdariffa* L

**DOI:** 10.64898/2026.07.27.741135

**Authors:** Branly Wilfrid Effa Effa, Dick Sadler Demikoyo Kanghou, Stéphane Mibemu Guibinga, Stevyna Mboumba-Mougola, Axelle Kedine Bindamba, Dieudonné Ndzengboro Endamane, Yves Anaclet Bagafou, Pamphile Nguema Ndoutoumou

**Affiliations:** Laboratoire de Biotechnologies Végétales, Institut de Recherches Agronomiques et Forestières (IRAF), Centre National de la Recherche Scientifique et Technologique (CENAREST) BP 2246; Laboratoire de Pédologie, Institut de Recherches Agronomiques et Forestières (IRAF), Centre National de la Recherche Scientifique et Technologique (CENAREST) BP 2246

**Keywords:** *Hibiscus sabdariffa*, growth, development, black soil, potting soil, treatments

## Abstract

*Hibiscus sabdariffa* L. var. *sabdariffa* is a plant prized in many tropical countries for its leaves and calyxes. Consequently, *H. sabdariffa* plays an important socio-economic role for many communities. Our study aimed to assess the effect of different substrates on the growth and development of *H. sabdariffa*. Three combinations were created by mixing black soil with potting soil, resulting in three substrates or treatments in addition to the control treatment (black soil). Each seeded substrate had eight replicates, resulting in 32 experimental units, arranged in a completely randomised design. The treatments were composed as follows: T0 (100% black soil, 0% potting soil), T1 (75% black soil, 25% potting soil), T2 (50% black soil, 50% potting soil) and T3 (25% black soil, 75% potting soil). The parameters measured were as follows: plant height, plant diameter, number of leaves per plant, leaf area of the plants and fresh above-ground biomass of the plants. The results showed that treatment T0 resulted in better growth and development of *H. sabdariffa*.

## Introduction

The sedentarization of humans led to the selection of plants through domestication, primarily for food and medicinal purposes. The hunter-gatherers of more than 10,000 years ago (in the Neolithic) are the initiators of domestication and the pioneers of agriculture (Diamond 2002). Humans need a healthy, balanced diet of plant and animal based foods to stay healthy. Among the plants, there is a plant called Roselle (*Hibiscus sabdariffa* L.) belongs to the Malvaceae family (Li et al. 2022; Mouhi et al. 2026; Sreenivasan et al. 2026). *H. sabdariffa*, a flowering plant of the genus *Hibiscus*, is native to Africa, probably West Africa (Gomaa et al. 2025). There are mainly two varieties of *H. sabdariffa*, the variety *H. sabdariffa cv sabdariffa* and the variety *H. sabdariffa cv altissima* which is rich in fiber (M’be et al. 2023). It spread to Asia and the Caribbean during the 16^th^ and 17^th^ centuries (Gomaa et al. 2025). However, *H. sabdariffa* is mainly cultivated in tropical and subtropical regions, particularly in South Asia and Africa (Paramadhas et al. 2026). *H. sabdariffa* is a herbaceous or sub-shrub plant, annual or perennial, woody base, up to 2 to 3.5 m high. Its cylindrical stems, generally red, are tender to almost tender (Mohamed et al. 2007; Ahmed et Khalid 2024). Its leaves, composed of 3 to 5 lobes, are 8 to 15 cm long. The flower, 8-10 cm in diameter, is white to pale yellow in colour, with a dark red mark at the base of each petal, the calyx is 1 to 2 cm wide at its base and the ripe, fleshy and bright red fruits are 3 to 3.5 cm long (Gomaa et al. 2025). Among the organs of *H. sabdariffa*, the fleshy red calyxes are the most prized. So far, only two types of *H. sabdariffa* calyxes have been described, one red and the other white, whose composition is similar except for the anthocyanins (M’be et al. 2023). When fresh, they are used in the preparation of several foods and beverages such as wine, juice, jams, jellies, syrups, gelatin, puddings, cakes, ice creams, and flavorings (Salami & Afolayan 2020). Dried, however, they are used, among other things, in the preparation of tea (Nguyen & Chuyen 2020; Paramadhas et al. 2026). The cultivation period of *H. sabdariffa* is about six months, from planting to harvest (Rajesh et al. 2020).

*H. sabdariffa* immediately attracted attention due to the many health benefits that have been reported (Juhari et al. 2018 ; Pattanittum et al. 2021). In traditional medicine, In rural areas of many developing countries, people use *H. sabdariffa* for medicinal purposes (Chikhoune et al. 2017; Mouhi et al. 2026). The red varieties of *H. sabdariffa* are used to promote cardiovascular health, prevent high blood pressure, fever and liver disorders, inhibit the growth of microorganisms, and act as a diuretic, digestive aid and sedative (Da-Costa-Rocha et *al*., 2014 ; Kokare et *al*., 2025). They also possess cyclooxygenase-inhibiting and antioxidant properties (M’be et al. 2023 ; Kokare et *al*., 2025). *H. sabdariffa* is also used in the formulation of certain cosmetic (Ismail et al. 2008 ; M’be et al. 2023 ; Paramadhas et al. 2026) and pharmaceutical products (Paramadhas et al. 2026). In sub-Saharan Africa, *H. sabdariffa* is of considerable economic importance (Majdoub et al. 2021). However, the main production areas in sub-Saharan Africa are Sudan, Senegal and Mali (Gomaa et al. 2025).

In Gabon, *H. sabdariffa* is mainly grown for food and medicinal purposes. However, the level of *H. sabdariffa* production in Gabon is insufficient to meet the local population’s needs for this commodity. The quality of the soil is one of the key factor for improving the quantity and quality of plant production, because plant yields are closely linked to the quality of the soil in which they are grown (Li et al. 2025). The chemical poverty of Gabonese soils has a negative impact on the growth and yields of *H. sabdariffa* (Ognalaga et *al*., 2015). It is therefore necessary to address this problem by finding alternative solutions. As the soil is the main source of mineral salts and water for the plants grown there, it therefore requires a little more attention in order to minimise the yield reductions observed in some smallholder plantations. Potting soils are routinely used in the greenhouses of research centres and institutes.

To this end, our study aims to evaluate the effect of three different substrates—comprising mixtures of soil and potting soil—on the growth and development of *H. sabdariffa*, compared with a substrate consisting solely of soil.

## Material and Method

The study was conducted at the “Institut de Recherches Agronomiques et Forestières” (0° 25’ 16, 571” N and 9° 52 25’ 55, 518” E).

### Plant material

In this study, we used the *H. sabdariffa* Koor variety with red calyxes (M’be et al., 2023), purchased from a agricultural shop in Libreville.

### Substrate composition

The substrates used were prepared from mixtures of soil collected from the undergrowth at a 30cm depth (Munthali et al. 2025) and a potting soil composed of black peat, coconut fibers, plant compost and PG mix 14-16-18 fertilizer. These substrates were the following: T0 (100% soil, 0% potting soil), T1 (75% soil, 25% potting soil), T2 (50% soil, 50% potting soil) and T3 (25% soil, 75% potting soil). They were analyzed and the results of these analysis are summarized in the table 1 below.

**Table 1:** Chemical characteristics of the treatments.

| Chemical characteristics | Substrates |  |  |  |
| --- | --- | --- | --- | --- |
|  | T0 | T1 | T2 | T3 |
| PH water | 6,7 | 6,5 | 6,5 | 6,8 |
| PH<sub>KCL</sub> | 5,3 | 4,8 | 4,8 | 4,9 |
| Total organic Carbon (g/kg) | 14,8 | 92,3 | 169,8 | 247,3 |
| Total Nitrogen (mg/g) | 0,9 | 3,7 | 6,45 | 9,225 |
| C/N | 16 | 24,9 | 26,32 | 26,8 |

T0 = 100% soil, 0% potting soil ; T1= 75% soil, 25% potting soil ; T2 = 50% soil, 50% potting soil and T3 = 25% soil, 75% potting soil.

### Experimental procedures

The experiment was carried out under a shade net. Soil collected from the undergrowth was sieved through a 2-mm sieve (Ling et al. 2025 ; Zeng et al. 2025) to separate it from coarse plant and non-plant matter, then placed in two 20-litre containers covered with plastic film was disinfected with water heated to 100ºC, 24 hours before being mixed with the potting soil in the following proportions: 0 volume potting soil to 4 volumes soil; 1 volume potting soil to 3 volumes soil; 2 volumes potting soil to 2 volumes soil; 3 volumes potting soil to 1 volume soil (table). Each of the mixtures constituted a substrate, resulting in 4 substrates or treatments. Before sowing, each substrate was placed in 8 black polyethylene nursery bags measuring 20×26 cm and then placed under the shade net. Direct sowing was carried out using two *H. sabdariffa* seeds per pot, with one seed per hole. Two holes, approximately 1 cm deep and spaced 1 cm apart, were dug in the centre of each pot, and one seed was placed in the bottom of each hole. Thinning was carried out to leave one vigorous plant per pot, two weeks after sowing . The trial lasted 45 days after sowing. Experiment was laid out in a Randomized Complete Block Design (RCBD) with 4 treatments of 8 replicates each. All the polyethylene bags containing the substrates were kept well watered throughout the trial.

### Parametres measured

In order to identify the appropriate substrate for the growth and development of *H. sabdariffa*, plant height, plant diameter, leaves number and leaf area were measured once a week in all the treatments. Height of the plants was measured using a retractable tape measure from the collar to the point where the last branch joins the stem; the diameter of the plants was measured using a manual caliper at the collar; the number of leaves on each plant was counted; The leaf area of each plant was determined by taking the mean of the leaf areas of the three most representative leaves on that plant; the fresh biomass of the shoot parts of each plant was measured at the end of the experiment using a precision balance.

### Statistical analysis

Data and statistical analysis were performed using the software Rstudio, means for each parameter were compared between treatments using ANOVA and Fisher LSD tests at 5%. (Fisher 1937). Means were considered to be significantly different when the difference between treatments was greater than the LSD value calculated from the ANOVA. This is indicated in the figure by asterisks; if there is more than one asterisk, the difference is highly significant.

## Result

### Plant height

Analysis of variance for plant height revealed a highly significant difference between the treatments (Figure 1) with *p-value < 0*.*0001= 7*.*1e-06*. The highest average height was obtained in the control treatment T0 (32.67 cm), followed by treatments T1 (23.72 cm), T2 (18.62 cm) and T3 (16.67 cm).

**Figure 1.**
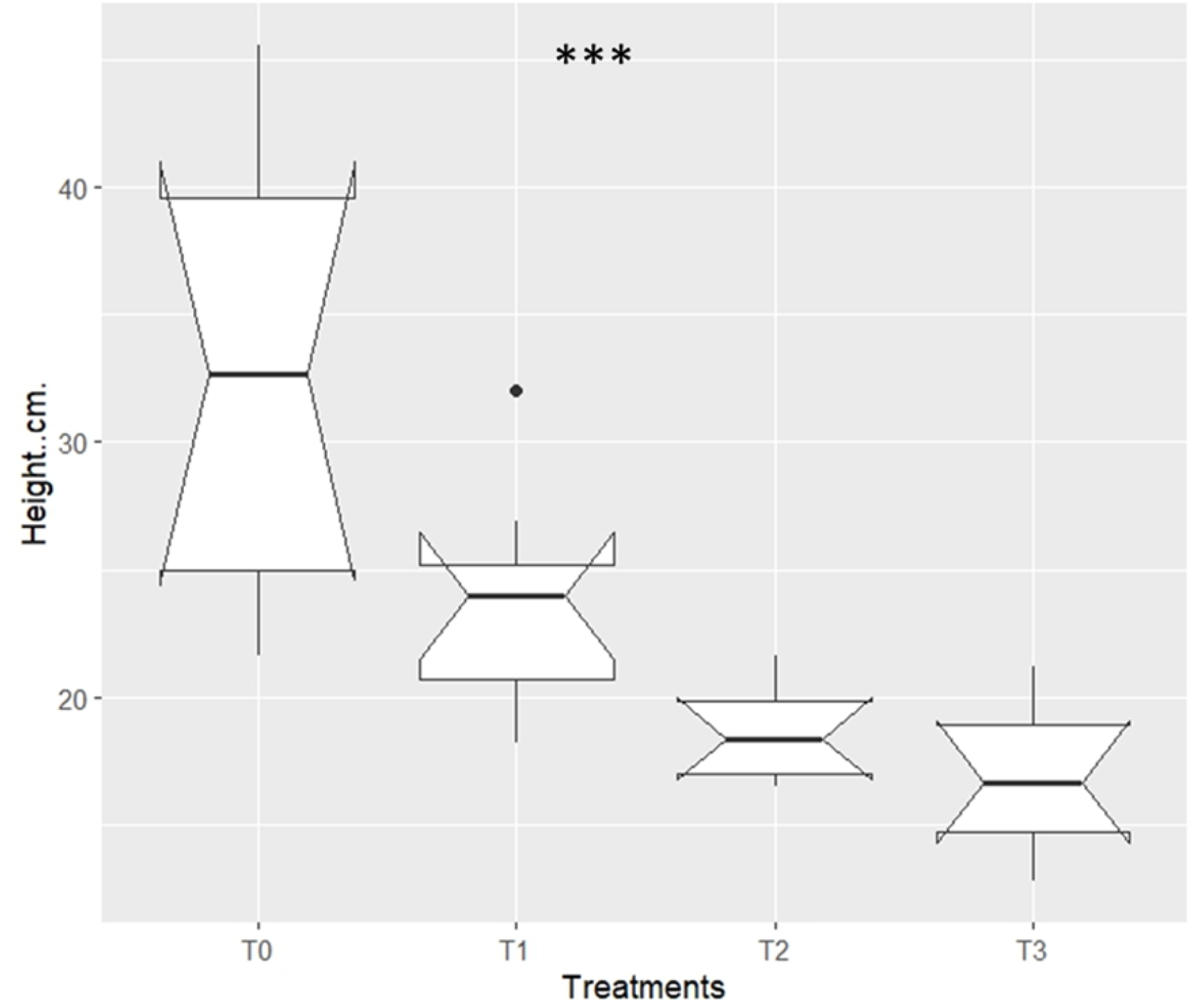
Plant height 45 days after sowing. T0 =100% soil ; T1 = 75% soil and 25% potting soil ; T2 = 50% soil and 50% potting soil ; T3 = 25% soil and 75% potting soil. n=8, ***: *p-value < 0*.*0001*.

### Shoot diameter

Analysis of variance for seedling diameter revealed a highly significant difference between the treatments (Figure 2) with *p-value < 0*.*001*. The highest mean seedling diameter was 3.33 mm, obtained in treatment T0. The mean seedling diameters obtained in treatments T1, T2 and T3 were 2.49 mm, 2.47 mm and 2.33 mm respectively.

**Figure 2.**
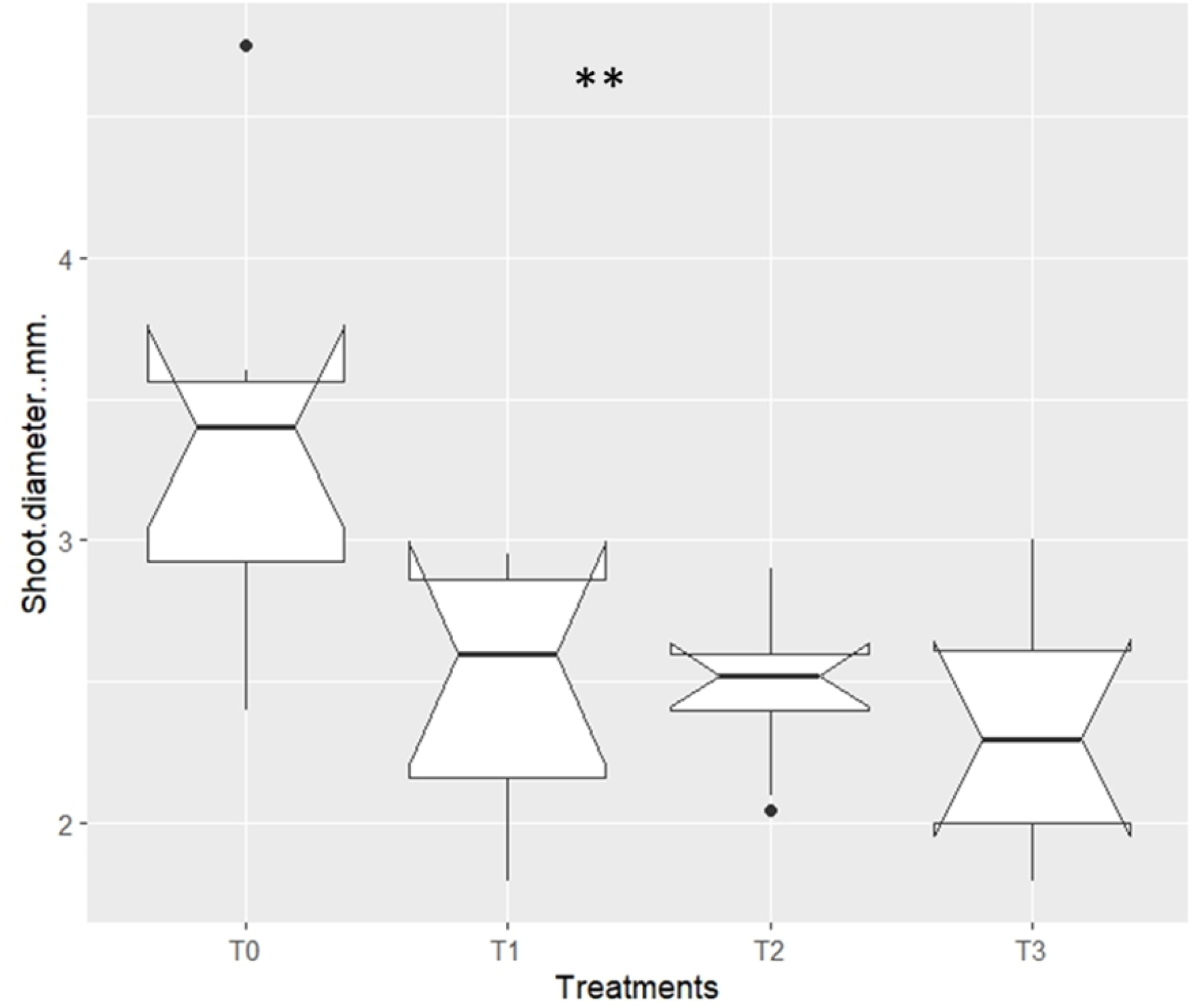
Plant diameter 45 days after sowing. T0 =100% soil ; T1 = 75% soil and 25% potting soil ; T2 = 50% soil and 50% potting soil ; T3 = 25% soil and 75% potting soil. n=8, **: *p-value < 0*.*001*.

### Leaves number

Analysis of variance for the number of leaves per plant revealed a highly significant difference between the treatments (Figure 3) with *p-value < 0*.*0001*. The highest number of leaves was 9.87, recorded in treatment T0. The lowest number of leaves was 6, recorded in treatment T3.

**Figure 3.**
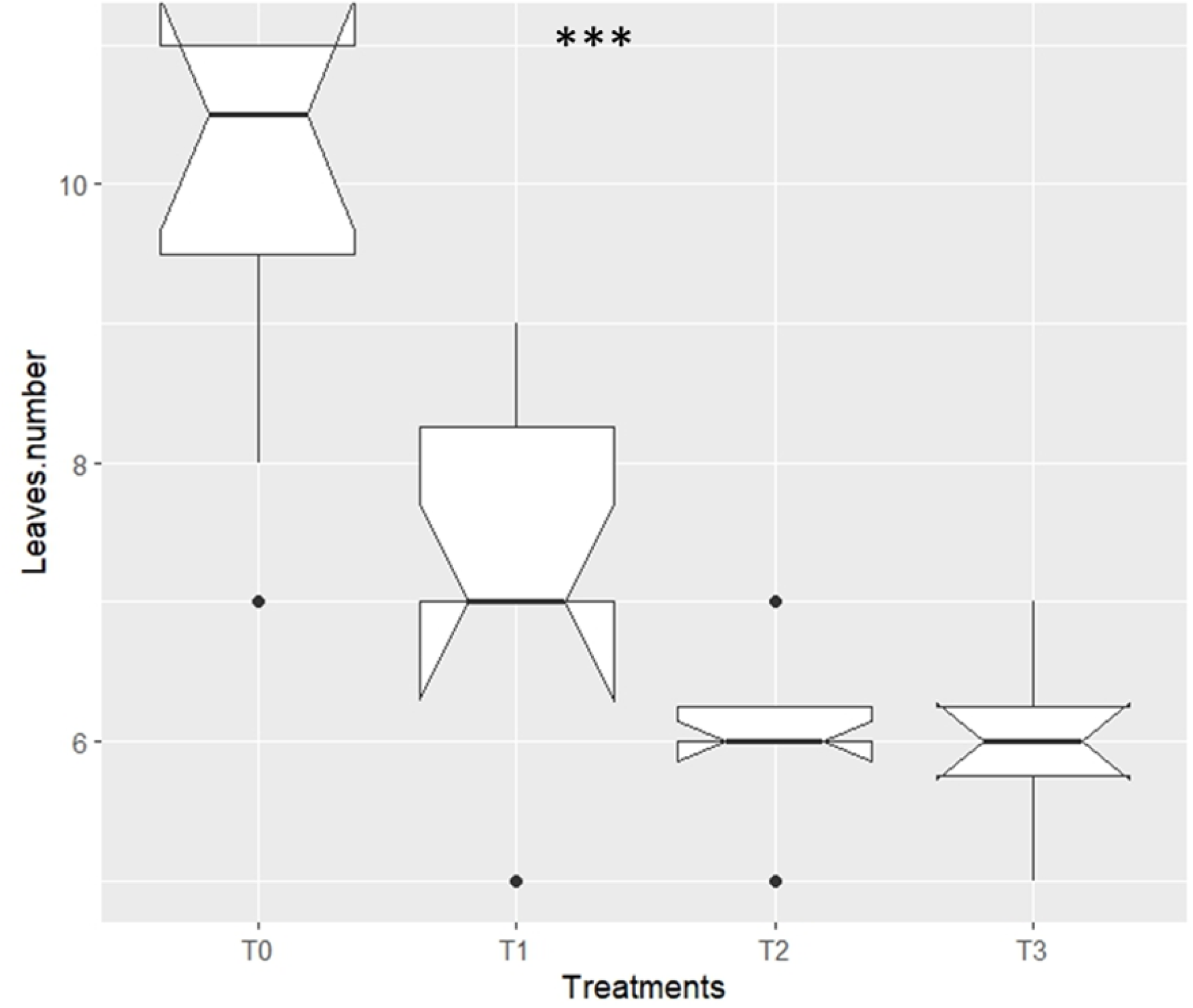
Plant diameter 45 days after sowing. T0 =100% soil ; T1 = 75% soil and 25% potting soil ; T2 = 50% soil and 50% potting soil ; T3 = 25% soil and 75% potting soil. n=8, ***: *p-value < 0*.*0001*.

### Leaf area

Analysis of variance for leaf area revealed a highly significant difference between the treatments (Figure 4) with *p-value < 0*.*0001*. The maximum value was 70.58 cm^2^, recorded for treatment T0. The values recorded for the other treatments were 31.75 cm^2^ for T1, 16.88 cm^2^ for T2 and 15.84 cm^2^ for T3.

**Figure 4.**
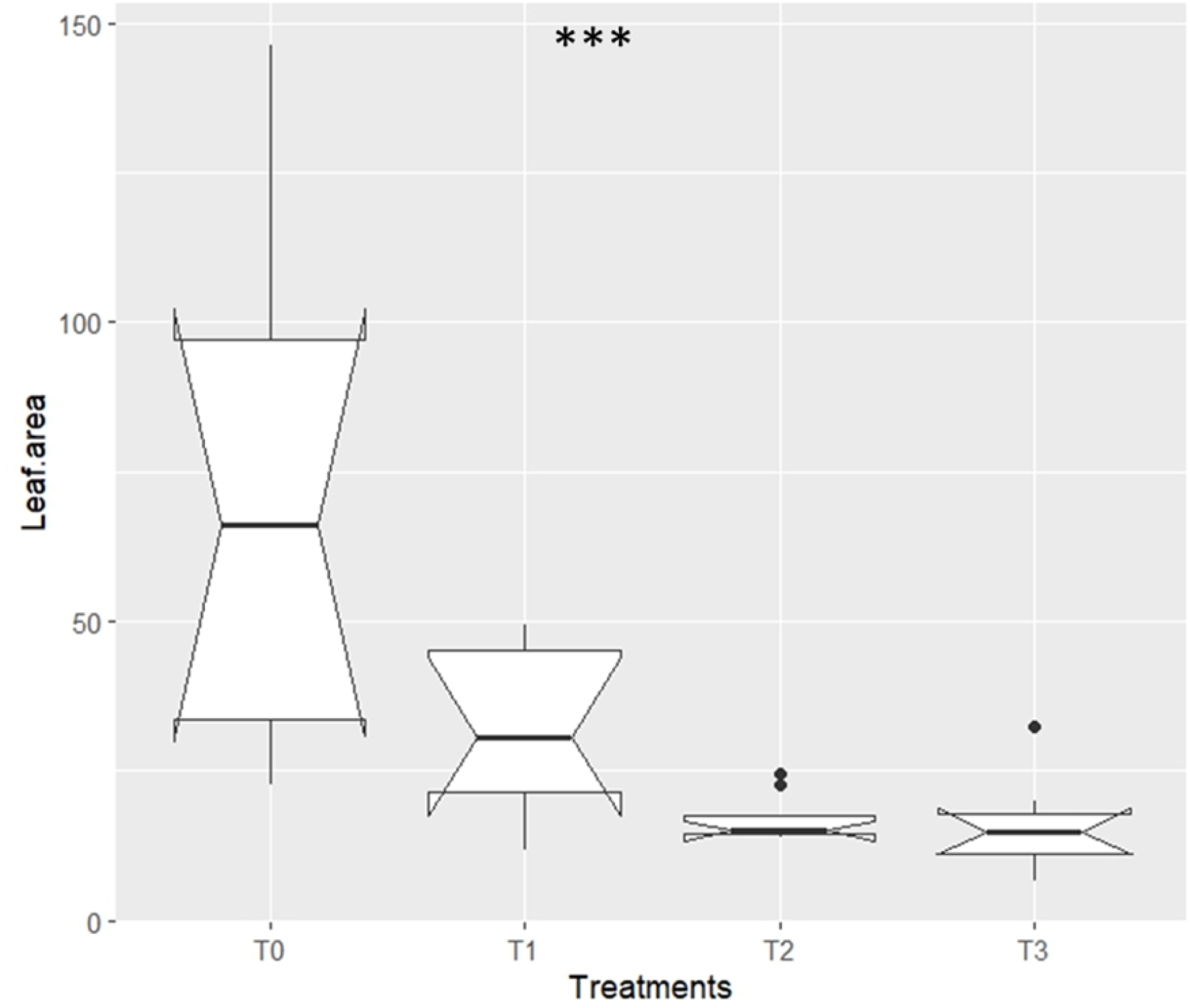
Leaf aera 45 days after sowing. T0 =100% soil ; T1 = 75% soil and 25% potting soil ; T2 = 50% soil and 50% potting soil ; T3 = 25% soil and 75% potting soil. n=8, ***: *p-value < 0*.*0001*.

### Shoot fresh biomass

Analysis of variance for shoot fresh biomass revealed a highly significant difference between treatments (Figure 5) with *p-value < 0*.*0001*. The highest shoot fresh biomass was obtained by treatment T0, with a value of 5.77 g. Treatments T1, T2 and T3 yielded the lowest shoot fresh biomasses, with 2.12 g, 1.17 g and 1 g respectively.

**Figure 5.**
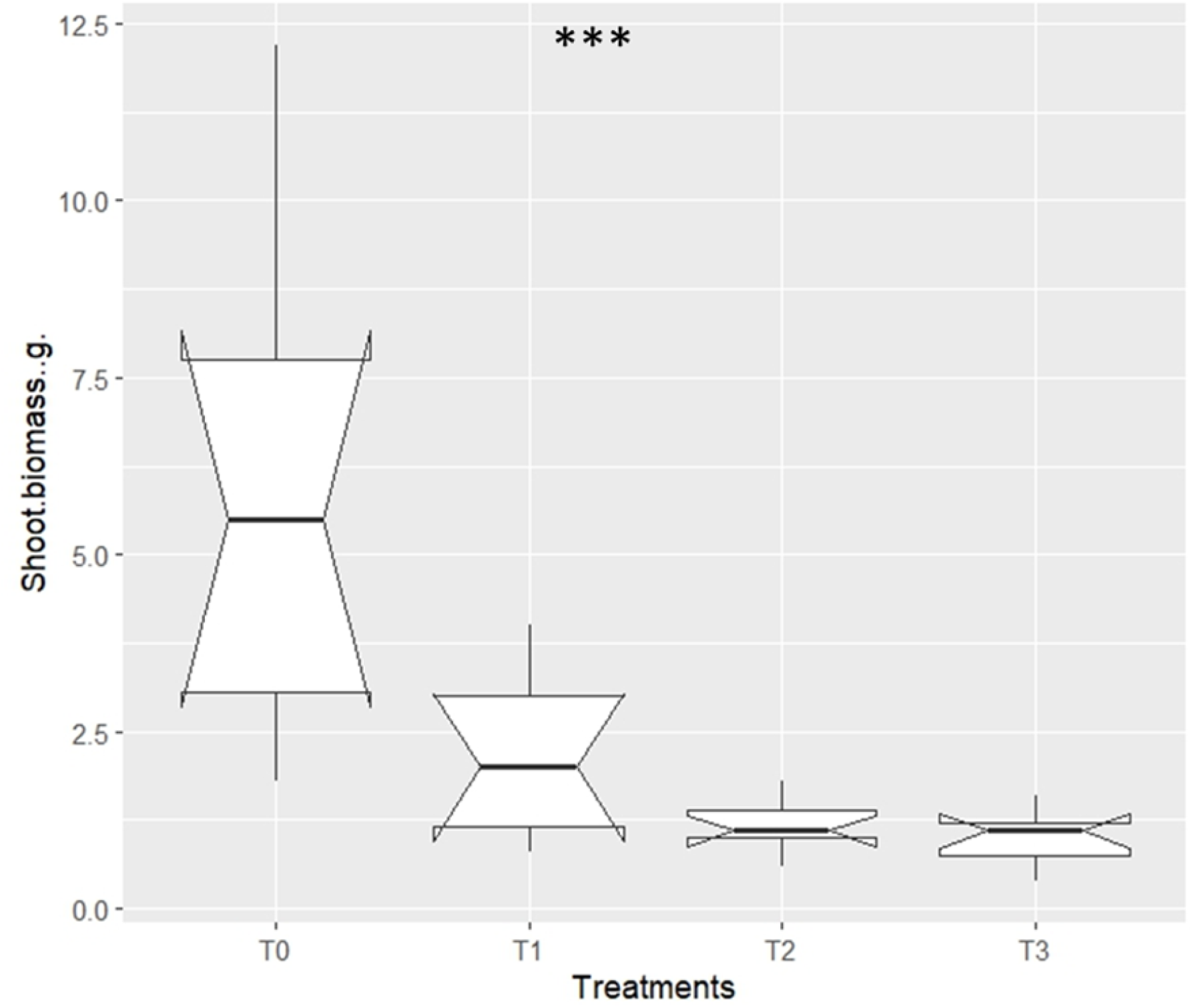
Shoot fresh biomass 45 days after sowing. T0 =100% soil ; T1 = 75% soil and 25% potting soil ; T2 = 50% soil and 50% potting soil ; T3 = 25% soil and 75% potting soil. n=8, ***: *p-value < 0*.*0001*.

### Correlation between parameters

Analysis of the correlation between the various parameters assessed revealed that there is a very strong correlation between all the parameters, with an excellent correlation (9.9) between shoot biomass and leaf area (Figure 6).

**Figure 6.**
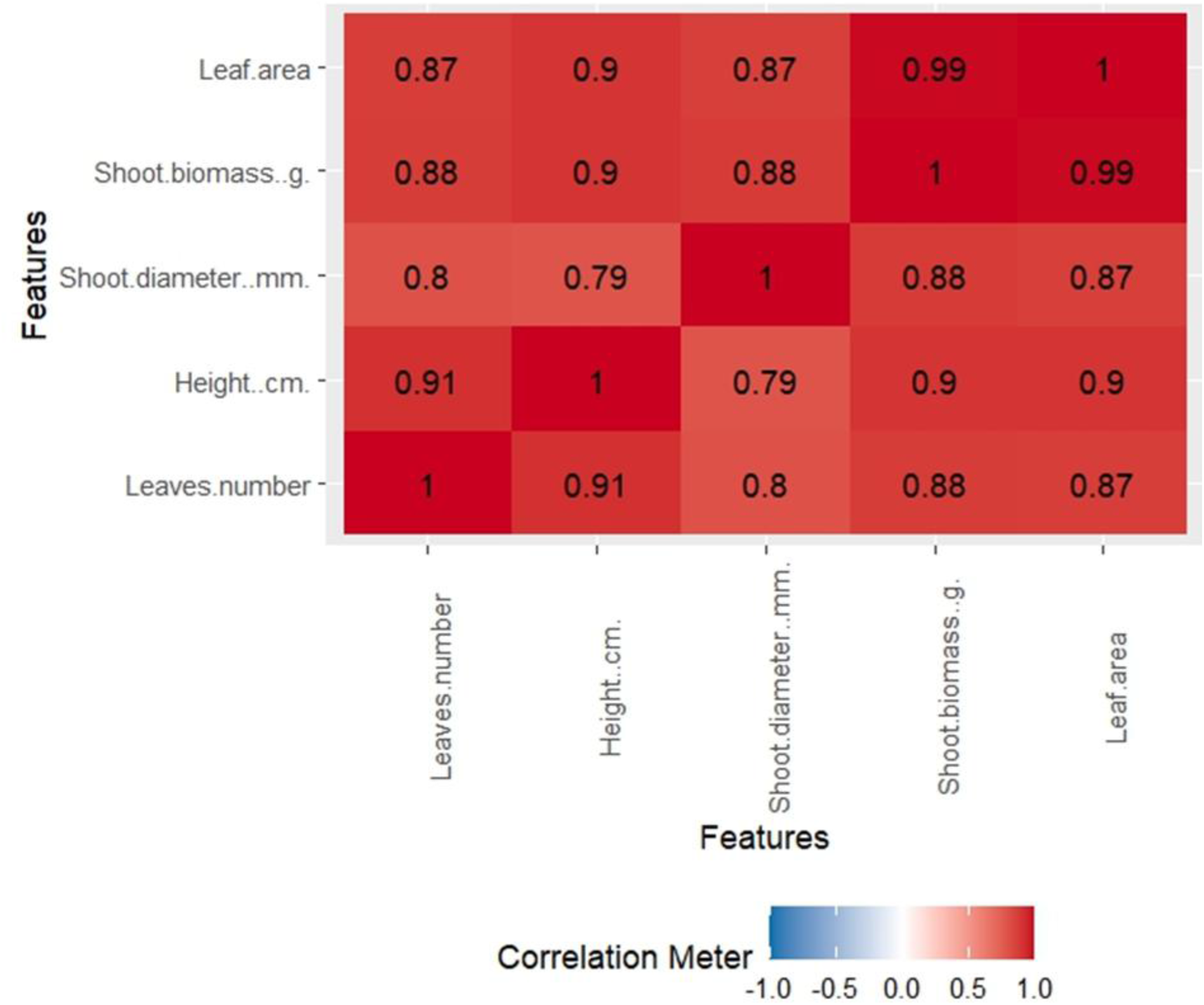
Relationship between parameters depending on treatments, n=8.

## Discussion

Although there is a very good correlation between the parameters observed independently of the treatments, results shows that T0 treatment offers better growth and development conditions for *H. sabdariffa*. Thus, growth and development conditions of *H. sabdariffa* become increasingly less favorable as the amount of potting soil increases and the amount of soil decreases. This led us to look at soil C/N ratio, which refers to the mass ratio between the total organic carbon in that soil and the total nitrogen; it is one of the most reliable indicators of soil quality (Zinn et al. 2018). Therefore, C/N ratio is respectively 16, 24.9, 26.32, 26.8 for T0, T1, T2 and T3. Growth and development of *H. sabdariffa* plants being greater with treatment T0, whose C/N ratio is 16, would offer better conditions for growth and development to *H. sabdariffa*. Indeed, a C/N ratio between 1 and 20 allows for rapid mineralization and nitrogen availability to plants (Brust 2019), contributing to the healthy growth and development of plants such as *H. sabdariffa*. The potting soil used in our experiment did not provide conditions for good plant growth and development due to its high C/N ratio. This ratio indicates reduced N availability, however N is an essential element for plants as it is a structural component of chlorophyll, enzymes such as Ribulose-1,5-bisphosphate carboxylase/oxygenase (Rubisco), and is crucial for photosynthesis (Chakrawal et al. 2026). Rubisco is crucial for photosynthesis and for plant growth and development (Yamori et al. 2026), N unavailability in soil would lead to reduced growth and development in plants cultivated there; this is what was observed in our study with *H. sabdariffa*.

## Conclusion

The results showed that for all the parameters studied, the highest values were obtained with treatment T0 (100% soil), with values more than double those of T1 and four times those of T2 and T3 for parameters such as leaf area and aboveground fresh biomass of each plant.

This work demonstrated that this potting soil, used as a substrate for container crops by some small-scale producers, is not suitable for plant growth and development.

